# Small variations in temperature and photoperiod under field conditions in South America modified spring barley phenology and foliar development

**DOI:** 10.64898/2026.08.27.747666

**Authors:** Nicolás Mastandrea, Gastón Quero, Luis Viega, Ariel Castro

## Abstract

**Context:** Barley production requires advanced knowledge of its response to changing environmental conditions in order to keep it competitive and sustainable.

**Aim:** Advance in the understanding of barley phenology and foliar development under South American field conditions.

**Methods:** 8 spring barley genotypes with differential phenology were studied in four field experiments under different temperature (through years and sowing dates) and photoperiod (through sowing dates) conditions. Time to anthesis, emergence to onset of stem elongation, stem elongation to anthesis, photoperiod response (PR) in these three traits, number of final leaves at anthesis (FLN) and phyllochron were measured.

**Key Results:** Time to anthesis and its subphases were shorter in late plantings but under similar photoperiod, temperature increased them. Cultivars have differential responses but with magnitude interactions and not crossover ones. Cultivar effects defined PR with no interaction with year (temperature). Temperature and photoperiod affected FLN, phyllochron and their relationship with time to anthesis. Under the shorter photoperiod, FLN and phyllochron were negatively correlated, FLN was higher in the warmer year and positively correlated with time to anthesis while phyllochron was not affected by temperature and had no correlation with time to anthesis. Under longer photoperiod, phyllochron was higher in the warmer year and time to anthesis was positively correlated with both FLN and phyllochron.

**Conclusions:** Cultivar basal thermal requirements and PR were consistent under the different studied conditions. Changes in temperature and photoperiod affected the relationship between time to anthesis, FLN and phyllochron suggesting that, although the three traits are arithmetically related, environmental conditions affect their balance.

**Online summary text:** Understanding barleýs responses to environmental changes are key for adapting the crop to future scenarios. Changes in temperature and photoperiod modify the crop final number of leaves, their rate of appearance, the length of the crop cycle and the interactions between these traits, implying important variations in the crop development. This information allows an improvement of crop practices in order to achieve a better crop and provides targets for breeding.

## Introduction

Barley (Hordeum vulgare L.) is one of the most important winter crops in the Southern Cone of South America (FAOSTAT 2026). In Uruguay, malting barley is the second most widely grown winter crop in terms of area and grain production (DIEA 2025). Almost all the grain produced is used for malting, mostly for the export market. In order to maintain and increase its presence in the production system and to secure its sustainability, appropriate technology (through breeding and improved agronomical practices) must be provided in a continuous basis.

Understanding phenology, the appropriate adjustment and management of cycles (Castro et al., 2008), and the adequate combination of developmental subphases under the most favorable conditions, among other factors, are key to a cultivar’s adaptation to its environment (Slafer 2003). Due to the differential response of genotypes to factors such as temperature and photoperiod (Boyd et al. 2003), the crop has been able to adapt to the prevailing local conditions across a wide range of environments, even in areas far from the place of origin. In the last decades, due to the exposure to shifting environment and events such as drought or extreme temperatures, fine-tuning crop physiology becomes more relevant to minimize the impact in yields (Fernández-Calleja et al. 2021).

Photoperiod and temperature are the main environmental factors that modulate the adaptation in temperate cereals like wheat or barley (Ochagavía et al. 2017; Hass et al. 2019), and these factors, their interaction and each cultivar’s specific characteristics can modify the pre-anthesis cycle’s development (Slafer and Rawson 1996). Barley quantitatively responds to long days, so its development rates increase when the photoperiod is extended, with varietal differences in sensitivity to day length (Miralles et al. 2014). If vernalization requirements exist, the flowering time is completed or accelerated when the crop is exposed to low temperatures (Ochagavía et al. 2022), whereas when those requirements are not satisfied, anthesis is delayed and may even fail to flower. Genotypes without vernalization requirements are classified as having spring habits.

The barley cycle can be divided into two developmental periods: emergence-stem elongation, and stem elongation-anthesis. In the first phase, the number of leaf and spikelet primordia per spike is differentiated; in the second phase, the number of grains per spike are determined (Kirby and Appleyard 1981; Waddington et al. 1983). Increased temperatures affect cereals in different ways, accelerating the rate of development of crops as wheat and causing reduction in the subphases (Slafer and Rawson 1994). Temperature may also affect the sensitivity to photoperiod at different developmental sub-phases (Slafer and Rawson 1996). After heading the phase duration is controlled by temperature in barley (Miralles et al. 2021) and rising temperatures had effect on grain filling.

The final leaf number depends on the rate and duration of the leaf differentiation process at the meristematic apex, which ends with the beginning of floral initiation (Slafer 2015). The duration of the differentiation process can modify the number of leaves per stem and time to anthesis (Slafer and Rawson 1997; Boyd et al. 2003; Borras et al. 2009). The time from leaf primordium differentiation to flag leaf emergence and expansion is associated with the number of differentiated leaf primordia and leaf emergence rate, which is affected by genetic and environmental factors (Kernich et al. 1995; Wilhelm and McMaster 1995).

The leaf emergence rate or its inverse, the phyllochron (defined as the interval between the appearances of two successive leaves in the same stem), are used as a measure of leaf development in grasses (Frank and Bauer 1995; Whilhem and McMaster 1995; Hay and Ellis 1998). In barley, changes in the phyllochron have been associated with different sowing dates, because of the length and rate of change of the post-emergence photoperiod (Kernich et al. 1995; Miralles et al., 2007) and temperature (Slafer and Rawson 1995). For Abeledo et al. (2004), changes in phyllochron throughout crop development could be due to different effects of cardinal temperatures on leaf emergence.

The fact that the mechanisms controlling development in spring barley are not fixed during one stage of development but are influenced by temperature before and during each stage of development (Frank and Bauer 1997), contributes to explain that the cardinal temperatures for each event (leaf emergence, stem elongation) can be different (Slafer and Rawson 1995) and highlights the complexity of the effects that each environment can have.

This study aims to advance in understanding the phenology of barley under the environmental conditions of Uruguay, focusing on the number of developed leaves and phyllochron, and their relationship with the length of the different phenological phases. For this purpose, a set of genotypes with differences in phenology was analyzed in four field experiments, defined by contrasting sowing dates and years.

## Materials and Methods

### Plant Material

Eight spring two-row barley cultivars of different origins were used: INIA Ceibo (Uruguay), Norteña Carumbé (released in Uruguay, developed at NDSU, USA), Estanzuela Quebracho (released in Uruguay, developed at the University of Western Australia), Baronesse (Germany), Bowman (USA), Danuta (Germany), Logan (USA), and Prior (Australia). The cultivars were selected based on a previous characterization of their phenology and photoperiod sensitivity (Locatelli et al. 2013, 2022). Details on the origin and coancestry of the cultivars are shown in Supplementary Table S1.

### Phenotyping

The developmental events measured were seedling emergence (e), onset of stem elongation (se), and awn emergence. According to Zadoks et al. (1974), these phases correspond to stages Z1.0, Z3.1, and Z4.9 (Fig. 1a). Awn emergence was recorded when 50% of the plants in the plot had awns approximately 1 cm above the flag leaf sheath. The awn emergence (Z4.9) stage is considered synonymous with anthesis (a), as described by Castro et al. (2017). Phenological phases were defined as follows: emergence–onset of stem elongation (e–se), onset of stem elongation–anthesis (se–a), and emergence–anthesis (e–a) (Fig. 1a). The equations used to calculate thermal time, phase duration, leaf appearance rate, phyllochron, and photoperiod response are presented in Table 1.

**Fig. 1.**
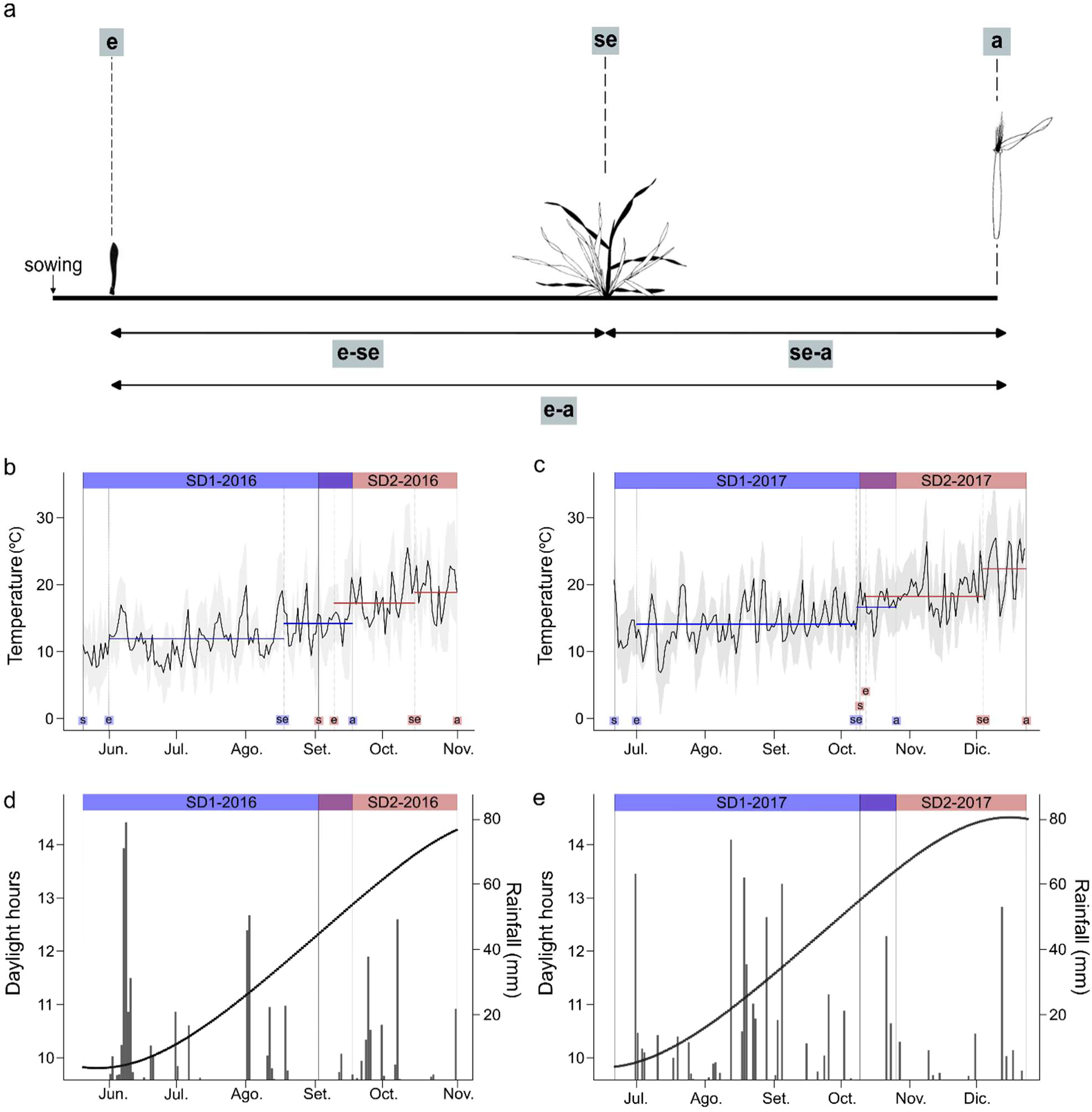
Schematic representation of barley developmental phases. **a)** Phenological stages adapted from Tottman et al. (1979). **b)** Mean temperature in 2016 since sowing date (s) and during the subphases emergence-onset of stem elongation (e-se), and onset of stem elongation-anthesis (se-a) for early sowing SD1 (blue) and late sowing SD2 (red). **c)** Mean temperature in 2017 since sowing date (s) and during the subphases emergence-onset of stem elongation (e-se) and onset of stem elongation-anthesis (se-a) for early sowing SD1 (blue) and late sowing SD2 (red). **d)** Photoperiod (line) and rainfall (bars) in 2016. **e)** Photoperiod (line) and rainfall (bars) in 2017.

**Table 1.** Equations for phenotyping.

| Trait | Equation | Num. |
| --- | --- | --- |
| Thermal time (°Cd) | $TT = \sum((T_{\max} - T_{\min})/2 - T_b)$ | Eqn 1 |
| Duration (e-se) | $\Delta TT_{(e-se)} = TT_{se} - TT_e$ | Eqn 2 |
| Duration (se-a) | $\Delta TT_{(se-a)} = TT_a - TT_{se}$ | Eqn 3 |
| Duration (e-a) | $\Delta TT_{(e-a)} = TT_a - TT_e$ | Eqn 4 |
| Leaf Appearance Rate<sup>†</sup> | $LAR = (N_{\text{leaf}} - \text{intercept}) / TT$ | Eqn 5 |
| Phyllochron | $Phy = 1/LAR$ | Eqn 6 |
| Photoperiod Response<sup>††</sup> | $PR(x) = \Delta TT_{(x \text{ SD1})} - \Delta TT_{(x \text{ SD2})}$ | Eqn 7 |

<sup>†</sup>To calculate the Leaf Appearance Rate (LAR), linear regressions were performed of leaf number (Nleaf) as a function of thermal time. Nleaf refers to the number of visible leaves.
<sup>††</sup>The subscript x in PR denotes any of the three phenological phases considered.

### Field experiments

Field experiments were conducted at the experimental field of the Facultad de Agronomía, Montevideo, Uruguay (34°51’S, 56°12’W). Four sowing dates were used: June 14th, 2016 (SD1-2016), June 27th, 2017 (SD1-2017), September 27th, 2016 (SD2-2016), and October 13th, 2017 (SD2-2017). The sowing dates were selected to correspond with photoperiods close to 10 h (SD1-2016 and SD1-2017) and greater than 12 h (SD2-2016 and SD2-2017). Planting was carried out manually in two row plots, each 1 m long, with 0.20 m between rows and 0.30 m between plots. A sowing density of 220 viable seeds m⁻¹ was used, and weeds were periodically removed manually. In each plot, five randomly chosen plants were monitored, and the number of leaves on the main stem (after the Z2.1 stage) was recorded, at least twice a week, according to Haun’s scale (Haun 1973) until reaching the flag leaf, considered the final leaf number (FLN). The environments determined by the different sowing dates in both years are described in (Fig. 1 b-e, Supplementary Tables S2-S3). Environmental data (rainfall, maximum and minimum air temperature) were recorded at an automatic station located in the same field.

### Statistical model analysis

All experiments were conducted using a randomized complete block design (RCBD), with three replications in the 2016 trials and four replications in the 2017 trials.

Each experiment was evaluated using the following linear mixed model (Eqn 8):

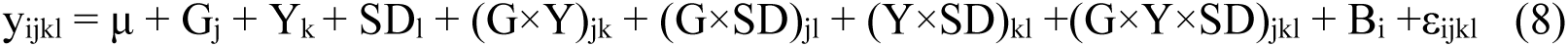

Where Y_ijkl_ is the response variable (i.e., ΔTT_(e-se),_PR_(x)_, μ is the overall mean, G_j_ is the fixed effect of genotype, Y_k_ is the fixed effect of year, SD_l_ is the fixed effect of sowing date. Interaction terms represent combined effects among genotype, year, and sowing date, B_i_ is the random effect of block, and ε_ijkl_ is the residual error. An ANOVA was performed to determine the significance of the main effects and their interactions, on the different response variables. Adjusted means for each treatment were obtained using the mixed model, in which genotype and environment factors were considered fixed, and blocks were considered random. The models and ANOVAs were fitted using the *lme4* package (Bates et al. 2015) in R (R Core Team 2023). Adjusted means and contrasts were obtained with the *emmeans* package (Lenth 2026).

## Results

### Duration of barley developmental stages

The fixed effects of the mixed model explained between 82 and 93% of the observed variance of the length of the different phases measured (Supplementary Table S4). Year and sowing date (SD) had significant effects on the duration of all the stages analyzed, whereas cultivar was significant for se–a and e–a. SD accounted for most of the observed variance, with 80, 52, and 78% for e–se, se–a, and e–a, respectively (Supplementary Table S5). As expected, SD1 had higher values for all the measured phenological phases compared with the late plantings. Year and cultivar, which was significant for se-a, and e-a, explained less than 10% of the observed variance, except in se–a, where cultivar explained nearly 28%. The warmer year (2017 had higher values for all the phenological traits in both planting dates. SD by Cultivar interaction was significant for all phases, while the three-way interaction was non-significant in all the stages analyzed (Supplementary Table S5). Duration of all stages was shorter in the second sowing date (SD2) in both years (Fig. 2).

**Fig. 2.**
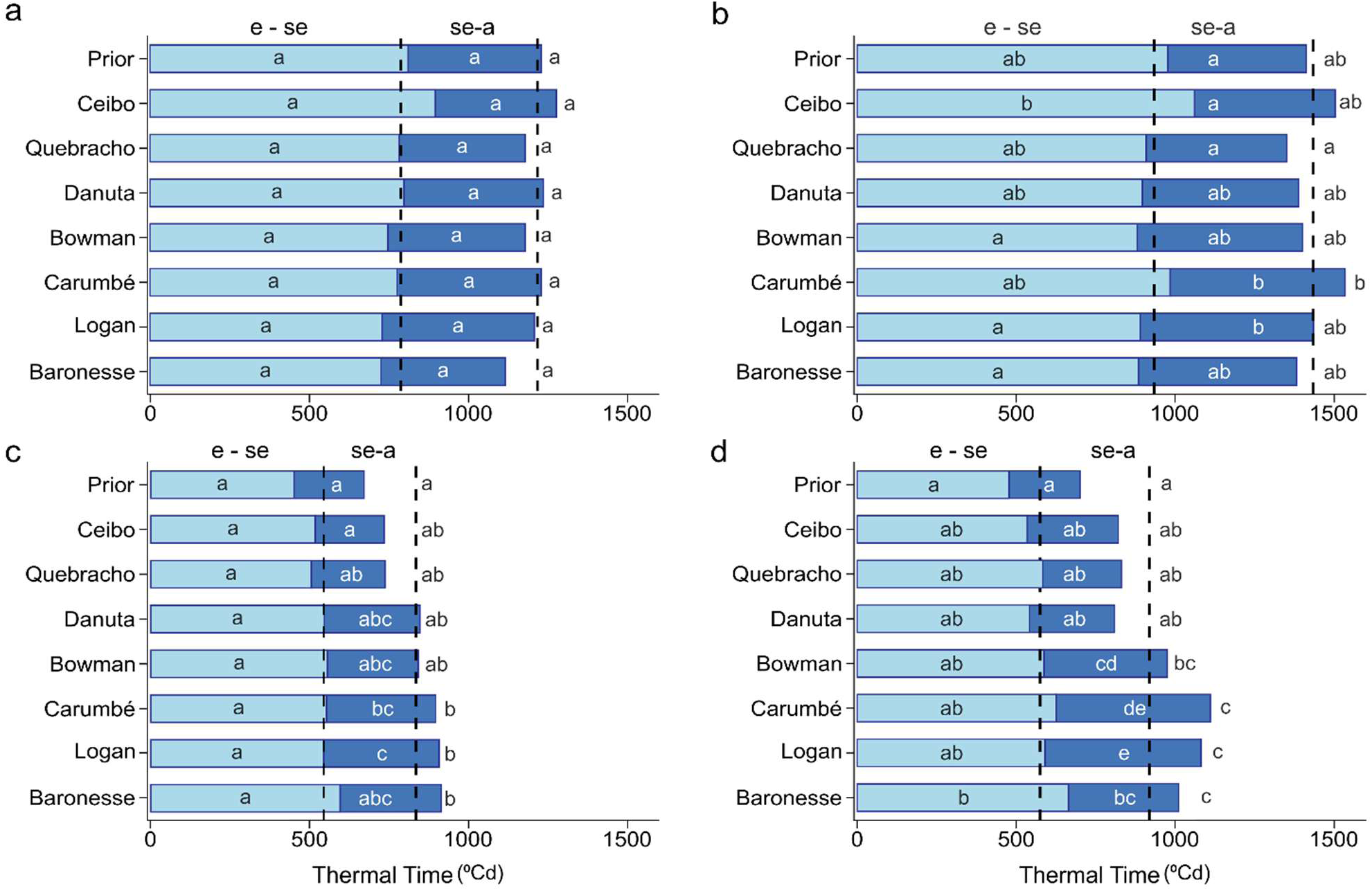
Thermal Time from emergence to anthesis (°Cd) and the subphases emergence–onset of stem elongation (e–se) and onset of stem elongation–anthesis (se–a) for eight genotypes across four environments: **a)** SD1-2016, **b)** SD1-2017, **c)** SD2-2016, and **d)** SD2-2017. Vertical dotted lines indicate the mean duration of e-se and e-a within each environment. Light blue bars represent the duration of the e–se subphase, dark blue bars represent the duration of the se–a subphase, and the full bar corresponds to the total duration from emergence to anthesis (e–a). Different letters in the bars indicate statistically significant differences among genotypes within each environment. Different letters next to the bars indicate statistically significant differences for thermal time to anthesis (e-a) (p < 0.05).

The mean thermal requirement for anthesis (e–a) in 2016 was 1206 °Cd and 820 °Cd for SD1 and SD2, respectively. In SD1-2016 there were no significant differences among cultivars, whereas in SD2-2016 three response groups were identified: a high thermal requirement group comprising Carumbé, Logan, and Baronesse; an intermediate group including Ceibo, Quebracho, Danuta, and Bowman; and a low requirement group consisting of Prior (Fig. 2c, Supplementary Table S6). In 2017, the e-a mean was 1426 °Cd in SD1 and 918 °Cd in SD2. This year, significant differences among cultivars were detected in both SDs. In SD1-2017, we defined three response types: a high thermal requirement for Carumbé; an intermediate group including Prior, Ceibo, Danuta, Bowman, Logan, and Baronesse; and a low requirement for Quebracho (Fig. 2b, Supplementary Table S6). In SD2-2017, four response patterns were identified: a high thermal requirement group composed of Carumbé, Logan, and Baronesse; an intermediate-high requirement for Bowman; an intermediate-low group composed of Ceibo, Quebracho, and Danuta; and a low requirement for Prior (Fig. 2d, Supplementary Table S6).

In 2016, the e–se subphase was 780 °Cd in SD1 and 532 °Cd in SD2, with no significant differences among cultivars at either sowing date. In 2017, this subphase was 935 °Cd in SD1 and 575 °Cd in SD2, with significant differences among cultivars (Fig. 2, Supplementary Table S6). In SD1-2017, three types of response were identified. The cultivar requiring the highest TT to complete the subphase was Ceibo, significantly differing from the group with the lowest duration, composed of Bowman, Logan, and Baronesse. A third group, consisting of Prior, Quebracho, Danuta, and Carumbé, exhibited an intermediate response, with no significant differences from the other groups (Fig. 2b, Supplementary Table S6). In SD2-2017, the cultivar with the lowest duration was Prior while Baronesse showed the longest duration. The intermediate response group for e-se includes the rest of the cultivars (Fig. 2d, Supplementary Table S6)

Duration of se–a in 2016 was 425 °Cd in SD1 and 286 °Cd in SD2. In SD1-2016, no significant differences among cultivars were observed, whereas in SD2-2016 four response patterns were identified: a high thermal requirement group composed of Logan and Carumbé; an intermediate-high requirement group including Bowman, Danuta, and Baronesse; an intermediate-low requirement for Quebracho; and a low requirement group consisting of Prior and Ceibo (Fig. 2c, Supplementary Table S6). In 2017, se-a was 490 °Cd in SD1 and 344 °Cd in SD2, and significant differences between cultivars were detected in both SDs. In SD1-2017, we identified three types of response: a high thermal requirement group (Logan and Carumbé); an intermediate group (Danuta, Bowman, and Baronesse), and a low requirement group (Prior, Ceibo, and Quebracho) (Fig. 2b, Supplementary Table S6). In SD2-2017, four response groups were detected: a high thermal requirement group (Logan and Carumbé), an intermediate-high requirement group (Baronesse and Bowman), an intermediate-low requirement group (Quebracho, Danuta, and Ceibo) and a low requirement for Prior (Fig. 2d, Supplementary Table S6).

### Photoperiod Response

The fixed effects of the mixed model explained between 55% and 79% of the observed variance (Supplementary Table S4). The cultivar had a significant effect on PR across all three stages, accounting for more than 70% of the variance in each stage (Supplementary Table S7). Year showed a significant effect on PR during the e–a, and e–se stages, explaining approximately 10% and 20% of the variance, respectively. Mean values for these two stages were higher in the warmer year. In contrast, the cultivar × year interaction did not have a significant effect on PR in any of the stages analyzed (Supplementary Table S7).

PR by cultivar had a similar pattern for e-se and e-a. Prior and Ceibo had the highest PR in both stages, and Bowman, Carumbé, Logan and Baronesse have the lower PR values (Fig. 3, Supplementary Table S8). Quebracho and Danuta had intermediate PR. Regarding se-a, cultivar differences were smaller and only Prior (high PR) had significant differences with Carumbé and Logan (low PR).

**Fig. 3.**
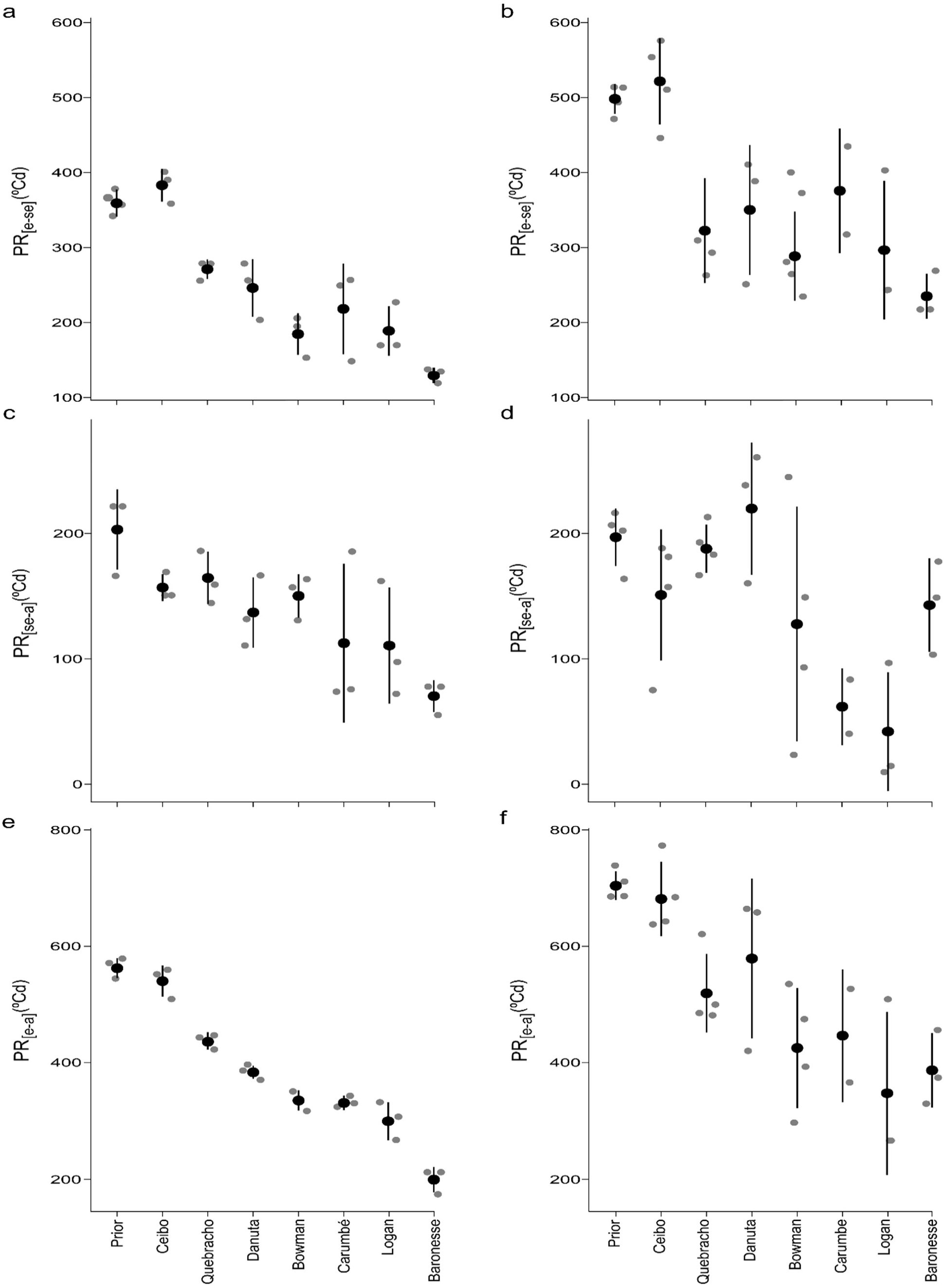
Photoperiod response (PR), expressed in thermal time (°Cd), across cultivars for time to anthesis (e–a) and its subphases e–se and se–a. Panels (a–b) show PR [e–se]; panels (c–d) show PR [se–a]; panels (e–f) show overall PR[e–a] by cultivar. Panels (a, c, e) correspond to 2016 and panels (b, d, f) to 2017.

### Final Leaf Number and Phyllochron

For FLN, the only non–significant source of variation was the three–way interaction cultivar × year × SD. Of the variance attributable to the fixed effects, SD was the dominant source, explaining 75% (Supplementary Table S9), with more leaves developed in SD1 than in SD2 (Table 2). Regarding the effect of the year, more leaves were developed in SD1 on the warmer year (2017), but for SD2 the values were similar in both years (Table 2).

**Table 2.** Final leaf number for all cultivars in each sowing date during the two years.

| Sowing Date | Cultivar | Year |  | Cultivar means |
| --- | --- | --- | --- | --- |
|  |  | 2016 | 2017 |  |
| SD1 | Prior | 11.4 b | 12.8 d | 12.1 |
|  | Ceibo | 11.2 b | 12.5 cd | 11.8 |
|  | Quebracho | 10.5 ab | 10.9 a | 10.7 |
|  | Danuta | 10.6 ab | 11.4 ab | 11.0 |
|  | Bowman | 9.8 a | 10.6 a | 10.2 |
|  | Carumbé | 11.6 b | 12.3 bcd | 11.9 |
|  | Logan | 9.7 a | 11.5 abc | 10.6 |
|  | Baronesse | 10.6 ab | 12.3 bcd | 11.4 |
|  |  | 10.7 A | 11.8 B | 11.2 |
| SD2 | Prior | 8.3 abc | 8.5 a | 8.4 |
|  | Ceibo | 8.0 ab | 8.3 a | 8.1 |
|  | Quebracho | 8.6 abc | 8.8 ab | 8.7 |
|  | Danuta | 8.5 abc | 8.7 ab | 8.6 |
|  | Bowman | 8.5 abc | 9.1 ab | 8.8 |
|  | Carumbé | 9.5 c | 9.0 ab | 9.2 |
|  | Logan | 8.1 a | 8.9 ab | 8.5 |
|  | Baronesse | 9.3 bc | 9.8 b | 9.5 |
|  |  | 8.6 A | 8.8 B | 8.7 |

SD1, early sowing date. SD2, late sowing date

For phyllochron, the only non–significant interaction was cultivar × year, whereas the two–way interactions and the three–way interaction involving SD were significant (Supplementary Table S9). Cultivar, Year, and SD had significant effects on phyllochron, accounting for 25%, 13%, and 37% of the variance attributable to the fixed effects, respectively (Supplementary Table S9).

Phyllochron values were higher in SD1 than SD2, with more differences in 2016. In the warmer year, the differences between sowing dates were reduced (Table 3). Regarding cultivar values, in 2016, all the genotypes had higher phyllochron during SD1, in 2017 the trend was the same except for Bowman, Carumbé and Logan which showed similar phyllochron in both sowing dates (Table 3).

**Table 3.** Phyllochron for all cultivars in each sowing date during the two years. Units in thermal time (°Cd leaf^-1^)

| Sowing Date | Cultivar | Year |  | Cultivar means |
| --- | --- | --- | --- | --- |
|  |  | 2016 | 2017 |  |
| SD1 | Prior | 120.5 abc | 110.0 a | 115.2 |
|  | Ceibo | 123.3 abc | 121.7 ab | 122.5 |
|  | Quebracho | 117.3 abc | 119.7 ab | 118.5 |
|  | Danuta | 123.7 bc | 127.2 b | 125.4 |
|  | Bowman | 130.7 c | 128.7 b | 129.7 |
|  | Carumbé | 108.8 a | 115.9 ab | 112.3 |
|  | Logan | 130.2 c | 130.9 b | 130.5 |
|  | Baronesse | 113.1 ab | 123.8 ab | 118.4 |
|  |  | 120.9 A | 122.3 A | 121.6 |
| SD2 | Prior | 72.8 a | 96.1 a | 84.4 |
|  | Ceibo | 69.2 a | 98.6 a | 83.9 |
|  | Quebracho | 63.9 a | 95.4 a | 79.6 |
|  | Danuta | 94.6 b | 115.5 b | 105.0 |
|  | Bowman | 101.0 b | 127.9 b | 114.4 |
|  | Carumbé | 95.3 b | 117.3 bc | 106.3 |
|  | Logan | 101.9 b | 133.0 c | 117.4 |
|  | Baronesse | 102.3 b | 113.8 b | 108.1 |
|  |  | 87.6 A | 112.2 B | 99.9 |

SD1, early sowing date. SD2, late sowing date

FLN was positively correlated with e–a in most environments, only in SD1-2016 this correlation was not significant (Fig. 4a). The correlations were higher, for both planting dates, in 2017. On the other hand, the phyllochron correlation with e-a, had contrasting values according to the sowing date: not significant in SD1, and strong positive correlation in SD2, regardless of the year (Fig. 4b). Finally, FLN and phyllochron correlation was strongly affected by the sowing date: negative and significant in SD1 and very low for SD2, regardless of the year (Fig. 4c).

**Fig. 4.**
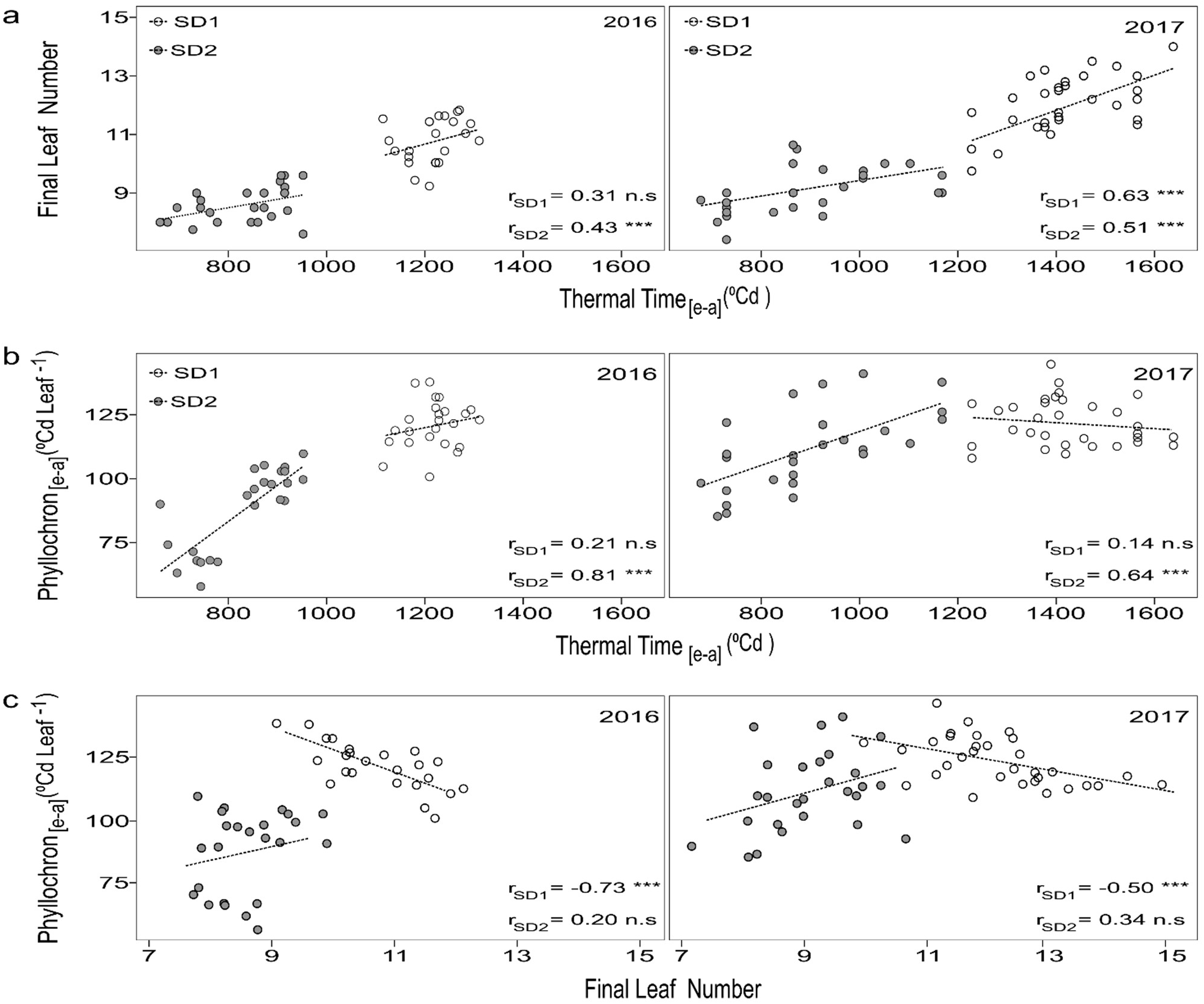
In panel a) relationship between Final Leaf Number and time to anthesis (Thermal Time_[e-a]_ (°Cd)); in panel b) relationship between Phyllochron_[e-a]_ (°Cd leaf^-1^) and time to anthesis (Thermal Time_[e-a]_ (°Cd)); in panel. c) Relationship between Phyllochron_[e-a]_ (°Cd leaf^-1^) and Final Leaf Number. In all cases early sowing (SD1) and late sowing (SD2) in 2016 and 2017 are represented. The Pearson correlation coefficient (r) is shown in each panel. *** indicates p < 0.05 (statistically significant correlation), n.s. indicates not statistically significant (p ≥ 0.05).

## Discussion

Barley is, as mentioned above, an important spring cash crop in the agricultural systems in the Southern Cone of South America. Worldwide, it has widely spread through a very diverse set of environments and established itself as a viable and useful commercial crop due, among other factors, to its high adaptability, which has produced consistently stable yields. Nevertheless, even in regions where it has been historically cultivated, changes in environmental conditions, even seemingly imperceptible ones, have affected its performance and threatened its sustainability. Deeper understanding of the responses of the crop to these small variations is required in order to develop the appropriate strategies (through breeding and agronomy) for better adaptation. In this research we use a limited set of genotypes, selected because of their contrasting phenology, in four experiments conducted under different temperature (through years and sowing dates) and photoperiod (through sowing dates) conditions to advance in the knowledge of barley phenology under South American environments. The number of cultivars used was limited by the available resources.

### Time to anthesis and photoperiod response

Both years, at early planting dates (SD1, associated with shorter days and average temperature close to 13°C) all cultivars had longer time to anthesis than in SD2, where all the genotypes reduced both pre-anthesis phases (e-se and se-a) (Fig. 2). This is consistent with the literature: in general, higher temperature and longer photoperiod has been associated with a reduction of the cycle (Boyd et al. 2003; Abeledo et al. 2004; Slafer et al. 2015; Parrado et al. 2023) in crops as wheat and barley. The differential magnitude of the reduction of the length of the e-a period and its phases between cultivars (measured by the PR) was within the expectations, with some cultivars (Prior and Ceibo in particular) having stronger reductions and others (Baronesse, Logan, Carumbé and Bowman) with much smaller ones. These results are consistent with previous studies under similar conditions (Locatelli et al. 2022). Of the four cultivars with lower PR, all but Baronesse originated in NDSU breeding program, and germplasm from that origin lack of photoperiod sensitivity. Baronesse has been reported also as lacking that sensitivity (Castro et al. 2017). The lack of genotype by year interaction for PR confirms that the trait is consistent between years and was not affected by the changes in temperature levels.

By comparing the results of similar sowing dates between years, we can analyze the effect of different mean temperatures under nearly equal photoperiods. The results showed that time to anthesis was longer in the warmer year, particularly in SD1. Although somewhat counter-intuitive and against the concept of higher temperature associated to reduced time to anthesis, it has been reported in the literature with several studies providing evidence that under certain conditions this may not occur. Rawson and Richards (1993) indicated that the time to anthesis is delayed by high temperatures in different photoperiods, Karsai et al. (2013), for spring barleys, showed that the duration until stem elongation and cycle is longer (in degree days) with increasing temperatures (from 13 to 23 °C). Ejaz and von Korff (2017) indicate that high-temperature environments (20 °C/16 °C vs. 28 °C/24 °C) delay flowering as there is repression, regardless of genotype, on the floral integrator FLOWERING LOCUS T1, which in barley is known as HvFT1 or Vrn-H3. Also, Hemming et al. (2012) report that in wheat and barley, the reproductive stage (double ridge) is delayed in short-day and high-temperatures scenarios; and Mastandrea et al. (2026) reported a delay in flowering time under warmer temperatures and short days for cultivars Ceibo and Carumbé, both used in this work. The novelty of our report derives from the fact that, unlike the cited findings, it was obtained from field experiments with narrow temperature differences between years. The other reports were obtained under high temperature and controlled conditions.

Slafer and Rawson (1996) reported that photoperiod sensitivity and optimum photoperiod change with temperature at each stage of development and this varies with genotype, so it could be considered that the delay in development generated in 2017 was a consequence of either the optimum temperature being exceeded or a negative effect of the warmer temperatures when days are short (Mastandrea et al., 2026). In general, optimum temperature is ignored because it is assumed that temperatures during growth are less than the optimal (Slafer and Rawson 1995).

### Final leaf number and phyllochron

FLN and phyllochron are partly defined by temperature and photoperiod, two factors that change in coordination (McMaster 2018). In this experiment both factors are defined through the combination of sowing date (affecting both) and year (affecting temperature). For all cultivars, the differences observed in both traits (FLN and phyllochron) between sowing dates were as expected for a quantitative long-day plant (Boyd et al. 2003), meaning fewer leaves developed in the main shoot and smaller phyllochron in late sowing dates compared with earlier ones (Table 2).

On the other side, the comparison between years provides a rather unexpected response: a higher FLN (in all cultivars) in SD1 for the warmest year. Hemming et al. (2012), as mentioned, detected a delay in the apex development at short photoperiods and temperatures of 25°C, and Mastandrea et al. (2026) described a slight increase in FLN in a warm environment (19 °C vs 12°C) under short day as potential cause of a reproductive development delay. As increasing temperature accelerates in a linear way the plastochron (McMaster 2018), it is possible that this effect was more noticeable on early sowing dates than in late ones. In the latter, although the temperature is higher (and also the differences between each yeaŕs temperature) the photoperiod is longer and limits the potential effect of temperature on FLN.

The phyllochron is considered stable (in degree days) during long periods of grass development (Rickman and Klepper 1995), although some authors consider a change during the development in crops like barley or wheat (Miralles et al. 2001; Brooking and Jamieson 2002; Abeledo et al. 2004). It has been reported a reducing effect of increased photoperiods on it (Miralles et al. 2007; Jamieson et al. 2008). In 2016 our results were consistent with that, with all cultivars presenting noticeable reductions (Table 3). But in the warmer year, the effect was smaller, with limited reduction and strong differences between cultivars, evidencing an interaction between photoperiod and temperature. The difference in temperature has no effect on phyllochron in SD1 (short photoperiod), but in late sowing dates (longer photoperiod) the warmer year had higher phyllochron. This is consistent with previous reports indicating higher phyllochron when temperature is rising under long photoperiods (Frank and Bauer 1997; Tamaki et al. 2002; Karsai et al. 2013), although the photoperiods from those studies were longer than in our experiments (16h, 14h and 16h respectively). According to Ochagavía et al. (2022), the variation is more affected by temperature than by genotypes. One possible explanation to this behavior is to have exceeded the optimal temperature for leaf emergence rate that has been described close to 20 °C for wheat and barley (Slafer and Rawson 1995, with 22°C for leaf emergence rate; Cao and Moss 1989, with 21°C for leaf emergence). In our experiments, this environment (long photoperiod, warmer year) had higher time to anthesis, but similar FLN, suggesting that phyllochron acted as a regulatory factor preventing more leaves. This is a novel fact, and more research is required to confirm it.

Under short photoperiod and low temperature (SD1-2016) FLN has no significant correlation with time to anthesis, while under warmer temperatures (SD1-2017) the association was significant and positive (Fig. 4) albeit with moderate correlation. Phyllochron showed no correlation with time to anthesis under short photoperiod, suggesting an independence from phenology in those conditions. The compensation between FLN and phyllochron was highlighted by their negative and significant correlation.

Under long photoperiod, both years, FLN and phyllochron had a significant and positive correlation with time to anthesis and no correlation between them, which suggests that under those conditions there was no “moderating” effect of phyllochron on FLN. The strong effect of photoperiod in reducing time to anthesis may be an explanation of this. Although particularly under short photoperiods FLN is associated with the duration of the vegetative stage (Pérez Gianmarco et al. 2018), due to a delayed floral induction and a rate of primordium initiation that is lesser affected (Parrado et al. 2023), the experiments shown the same association under long photoperiods, when the phases are reduced. Since daylength is considered the primary factor inducing plants to develop reproductive structures on the apex (Frank and Bauer 1995), probably in SD1, the cultivars expressed their full potential in leaf formation and development, but, in SD2 this association may be broken by higher photoperiod and temperatures.

The relationships between time to anthesis, FLN, and phyllochron summarized some aspects of the dynamics of development under contrasting conditions. The combination of temperature and photoperiod changed them, and our results evidenced that in some conditions they show independency from each other (although phyllochron is basically a ratio between the other two). In each condition (combinations of temperature and photoperiod) at least one of the three correlations was non-significant. In SD-2016 only FLN and phyllochron had a significant correlation, in SD1-2017 time to anthesis and phyllochron had no correlation and in SD2 (both years) FLN and phyllochron had no correlation. The last result evidenced some kind of independence of those two traits under longer photoperiods, which may be explained for differences in their genetic control in certain conditions. In order to further explore this hypothesis, a different approach is needed. We are presently working in a genetic mapping approach in order to advance in that direction.

Although we worked in only one location, it was representative (in terms of daylength and temperature) of the barley growing region in the southern cone of South America, and the combination of contrasting sowing dates and contrasting years in terms of temperature was adequate to study the effects of these factors in barley phenology under field conditions. Our results confirmed previous results and provided new and valuable information regarding the relationship between foliar development and phenology in this valuable crop.

## Conclusions

Temperature, photoperiod and genotype define barley phenology and our results, as expected, confirmed that. The relatively small differences in temperature and daylength between the environments in our study did not prevent the detection of effects (including interactions) in all the studied traits.

Cultivar responses to photoperiod for the length of the different phenological phases were significant and have low interaction with temperature. Temperature and photoperiod showed interaction with cultivar on the length of the different phases, but these interactions were mainly magnitude, not crossover ones. Good characterization regarding basal requirements and PR response can be used to obtain reasonable predictions of the genotype performance under the environmental scope of this study

Foliar development, measured by FLN and phyllochron, interacted with temperature and photoperiod. The changes detected in these traits under rising temperatures, highlight the need for further research in this area in order to fully estimate the possibilities of crop improvements through breeding or crop practices.

## Data availability

Data are available from the authors upon request.

## Declaration of funding (compulsory section)

This study was supported by projects of the Comisión Sectorial de Investigación Científica (CSIC), (financiación 1.1, programa 348) and funds form the Mesa Nacional de Entidades de Cebada Cervecera (MNECC) project, (Financiación 3.3)

## Acknowledgements (not compulsory)

The authors wish to thank the collaboration of Matias Antuoni with the establishment and maintenance of the field experiments, Yolanda Fernández for her invaluable assistance during the data collection, and also to undergraduate students from Facultad de Agronomía, UdelaR (Montevideo) who participated in the experiment activities. We also extend our acknowledgement to Maximiliano Verocai and Andrés Locatelli for their valuable suggestions to this manuscript.

## Supplementary material

Supplementary material can be accessed from the article page online.

