## Supplemental data for "Small variations in temperature and photoperiod under field conditions in South America modified spring barley phenology and foliar development"

**Supplementary material**

**Table S1:** Description of the germplasm set.

^†^The clustering is due to their coancestry, as reported by Locatelli et al. (2013).

| Cultivar | Country of origin. | Group^†^ |
| --- | --- | --- |
| Baronesse | Germany | A |
| Danuta | Germany | A |
| Prior | Australia | A |
| Estanzuela Quebracho | Australia | A |
| Bowman | USA | D |
| Logan | USA | D |
| Norteña Carumbé | Uruguay | D |
| INIA Ceibo/CLE 202 | Uruguay | E |

**Table S2:** Environmental conditions from emergence to onset of stem elongation (e–se).

| Environmental variable | SD1-2016 | SD2-2016 | SD1-2017 | SD2-2017 |
| --- | --- | --- | --- | --- |
| T. mean (°C) | 11.8 ± 2.9 | 17.2 ± 3.2 | 14.1 ± 3.1 | 18.1 ± 2.6 |
| T. max. media (°C) | 15.7 ± 4.1 | 22.1 ± 4.8 | 18.1 ± 3.8 | 23.0 ± 3.5 |
| T. min. media (°C) | 7.9 ± 2.7 | 12.3 ± 3.3 | 10.1 ± 3.5 | 13.1 ± 2.9 |
| Days with T > T.mean | 36 | 14 | 39 | 29 |
| Days with T> T.max.med. | 30 | 10 | 33 | 22 |
| Days with T< T.min.med. | 43 | 19 | 44 | 23 |
| Photoperiod (h) | 10.6 | 13.2 | 11.0 | 13.8 |
| Min. photoperiod (h) | 9.8 | 12.6 | 9.9 | 13.1 |
| Max. photoperiod (h) | 11.7 | 13.8 | 12.5 | 14.4 |
| Accumulated rainfall (mm) | 441.8 | 156.8 | 501.1 | 101.7 |

**Table S3:** Environmental conditions from emergence to onset of stem elongation (se–a).

| Environmental variable | SD1-2016 | SD2-2016 | SD1-2017 | SD2-2017 |
| --- | --- | --- | --- | --- |
| T. mean (°C) | 13.9 ± 2.3 | 19.1 ± 3.1 | 16.5 ± 2.2 | 22.1 ± 3.8 |
| T. max. media (°C) | 18.6 ± 3.9 | 24.8 ± 4.2 | 21.4 ± 3.2 | 27.4 ± 4.9 |
| T. min. media (°C) | 9.2 ± 2.5 | 13.4 ± 2.9 | 11.7 ± 2.9 | 16.8 ± 3.2 |
| Days with T > T.mean | 17 | 12 | 15 | 13 |
| Days with T> T.max.med. | 15 | 10 | 12 | 12 |
| Days with T< T.min.med. | 13 | 8 | 13 | 10 |
| Photoperiod (h) | 12.3 | 14.1 | 13.1 | 14.4 |
| Min. photoperiod (h) | 11.8 | 13.8 | 12.6 | 14.5 |
| Max. photoperiod (h) | 12.8 | 14.3 | 13.5 | 14.6 |
| Accumulated rainfall (mm) | 13.6 | 24.6 | 83.6 | 72.6 |

**Table S4.** Marginal and Conditional R² values for mixed models.

| **Trait** | **R² marginal** | **R² conditional** |
| --- | --- | --- |
| TT (e-se) | 0.82 | 0.852 |
| TT (se-a) | 0.84 | 0.839 |
| TT (e-a) | 0.93 | 0.931 |
| PR (e-se) | 0.79 | 0.88 |
| PR (se-a) | 0.55 | 0.55 |
| PR (e-a) | 0.76 | 0.89 |
| FLN | 0.66 | 0.66 |
| Phyllochron (e-a) | 0.62 | 0.62 |

**Table S5**: Type III ANOVA (Satterthwaite’s method) for subphases for thermal time.

| Trait | Fixed effects | Sum Sq | Mean Sq | NDF | DDF | F value | Pr(>F) | % Variance |
| --- | --- | --- | --- | --- | --- | --- | --- | --- |
| TT (e-se) | Cultivar | 60089 | 8584 | 7 | 72.9 | 1.50 | 0.18 | 2.01 |
|  | Year | 222751 | 222751 | 1 | 74.9 | 39.04 | ******* | 7.44 |
|  | Sowing date (SD) | 2398249 | 2398249 | 1 | 71.9 | 420.3 | ******* | **80.07** |
|  | Cultivar × Year | 15131 | 2162 | 7 | 71.8 | 0.38 | 0.91 | 0.51 |
|  | Cultivar × SD | 210925 | 30132 | 7 | 71.8 | 5.28 | ******* | 7.04 |
|  | Year × SD | 82782 | 82782 | 1 | 71.9 | 14.51 | ******* | 2.76 |
|  | Cultivar × Year × SD | 5173 | 739 | 7 | 71.8 | 0.13 | 0.99 | 0.17 |
| TT (se-a) | Cultivar | 288127 | 41161 | 7 | 75.0 | 20.96 | *** | 28.43 |
|  | Year | 100108 | 100108 | 1 | 75.0 | 50.98 | *** | 9.88 |
|  | Sowing date (SD) | 525175 | 525175 | 1 | 75.0 | 267.45 | *** | **51.83** |
|  | Cultivar × Year | 36701 | 5243 | 7 | 75.0 | 2.67 | * | 3.62 |
|  | Cultivar × SD | 46639 | 6663 | 7 | 75.0 | 3.393 | ** | 4.60 |
|  | Year × SD | 387 | 387 | 1 | 75.0 | 0.197 | 0.66 | 0.04 |
|  | Cultivar × Year × SD | 16153 | 2308 | 7 | 75.0 | 1.175 | 0.33 | 1.59 |
| TT (e-a) | Cultivar | 344843 | 49263 | 7 | 71.9 | 9.41 | *** | 5.18 |
|  | Year | 576568 | 576568 | 1 | 74.9 | 110.13 | *** | 8.66 |
|  | Sowing date (SD) | 5172797 | 5172797 | 1 | 71.9 | 988.05 | *** | **77.71** |
|  | Cultivar × Year | 76003 | 10858 | 7 | 71.8 | 2.07 | 0.06 | 1.14 |
|  | Cultivar × SD | 376162 | 53737 | 7 | 71.8 | 10.26 | *** | 5.65 |
|  | Year × SD | 95962 | 95962 | 1 | 71.9 | 18.33 | *** | 1.44 |
|  | Cultivar × Year × SD | 14122 | 2017 | 7 | 71.8 | 0.38 | 0.91 | 0.21 |

Significance codes: p < 0.01 ***; p < 0.01**; p < 0.05*

**Table S6:** Thermal time duration of subphases.

| Year | Sowing Dates | Genotypes | Subphases | | |
| --- | --- | --- | --- | --- | --- |
|  |  |  | e-se | se-a | e-a |
| 2016 | SD-1 | Prior | 805 a | 423 a | 1228 a |
|  |  | Ceibo | 895 a | 380 a | 1275 a |
|  |  | Quebracho | 777 a | 396 a | 1174 a |
|  |  | Danuta | 793 a | 438 a | 1231 a |
|  |  | Bowman | 742 a | 435 a | 1177 a |
|  |  | Carumbé | 771 a | 457 a | 1228 a |
|  |  | Logan | 732 a | 475 a | 1208 a |
|  |  | Baronesse | 723 a | 381 a | 1115 a |
|  | SD-2 | Prior | 445 a | 220 a | 666 a |
|  |  | Ceibo | 512 a | 223 a | 735 ab |
|  |  | Quebracho | 506 a | 232 ab | 738 ab |
|  |  | Danuta | 546 a | 301 abc | 848 ab |
|  |  | Bowman | 558 a | 285 abc | 842 ab |
|  |  | Carumbé | 552 a | 344 bc | 897 b |
|  |  | Logan | 543 a | 365 c | 908 b |
|  |  | Baronesse | 594 a | 321 abc | 915 b |
| 2017 | SD-1 | Prior | 979 ab | 437 a | 1416 ab |
|  |  | Ceibo | 1061 b | 441 a | 1503 ab |
|  |  | Quebracho | 909 ab | 442 a | 1350 a |
|  |  | Danuta | 896 ab | 492 ab | 1388 ab |
|  |  | Bowman | 880 a | 520 ab | 1400 ab |
|  |  | Carumbé | 984 ab | 550 b | 1534 b |
|  |  | Logan | 888 a | 546 b | 1434 ab |
|  |  | Baronesse | 885 a | 496 ab | 1381 ab |
|  | SD-2 | Prior | 477 a | 235 a | 712 a |
|  |  | Ceibo | 535 ab | 286 ab | 821 ab |
|  |  | Quebracho | 582 ab | 249 ab | 831 ab |
|  |  | Danuta | 542 ab | 267 ab | 809 ab |
|  |  | Bowman | 588 ab | 388 cd | 975 bc |
|  |  | Carumbé | 626 ab | 483 de | 1109 c |
|  |  | Logan | 588 ab | 499 e | 1087 c |
|  |  | Baronesse | 660 b | 349 bc | 1009 c |

The values are adjusted means, expressed in ºCd.

**Table S7:** Type III ANOVA (Satterthwaite’s method) for subphases for photoperiod response.

| Trait | Fixed effects | Sum Sq | Mean Sq | NDF | DDF | F value | Pr(>F) | % Variance |
| --- | --- | --- | --- | --- | --- | --- | --- | --- |
| PR  (e-se) | Cultivar | 4.5646 | 0.6521 | 7 | 31.8 | 35.25 | *** | 77.39 |
|  | Year | 1.1564 | 1.1564 | 1 | 33.4 | 62.50 | *** | 19.60 |
|  | Cultivar × Year | 0.1775 | 0.0254 | 7 | 31.8 | 1.37 | 0.252 | 3.01 |
| PR (se-a) | Cultivar | 87955 | 12565.0 | 7 | 35.0 | 6.37 | *** | 74.07 |
|  | Year | 737 | 737.4 | 1 | 35.0 | 0.37 | 0.545 | 0.62 |
|  | Cultivar × Year | 30055 | 4293.6 | 7 | 35.0 | 2.18 | 0.061 | 25.31 |
| PR  (e-a) | Cultivar | 734747 | 104964 | 7 | 31.84 | 40.34 | *** | 86.44 |
|  | Year | 88057 | 88057 | 1 | 33.04 | 33.84 | *** | 10.36 |
|  | Cultivar × Year | 27229 | 3890 | 7 | 31.84 | 1.49 | 0.205 | 3.20 |

Significance codes: p < 0.01 ***; p < 0.01**; p < 0.05*

**Table S8:** Photoperiod response across subphases.

| Year | Cultivar | Subphases | | |
| --- | --- | --- | --- | --- |
|  |  | PR_(e-se)_ | PR_(se-a)_ | PR_(e-a)_ |
| 2016 | Prior | 380 d | 203 b | 588 d |
|  | Ceibo | 405 d | 157 ab | 566 d |
|  | Quebracho | 287 cd | 165 ab | 462 cd |
|  | Danuta | 258 bc | 137 ab | 409 bc |
|  | Bowman | 194 ab | 150 ab | 361 bc |
|  | Carumbé | 224 bc | 113 ab | 357 abc |
|  | Logan | 198 b | 111 ab | 326 ab |
|  | Baronesse | 137 a | 70 a | 226 a |
|  |  | 260.4 *a* | 138.5 *a* | 411.8 *a* |
| 2017 | Prior | 502 cd | 202 c | 704 e |
|  | Ceibo | 523 d | 156 abc | 681 de |
|  | Quebracho | 321 ab | 192 bc | 519 bc |
|  | Danuta | 344 b | 224 c | 575 cd |
|  | Bowman | 288 ab | 132 abc | 425 ab |
|  | Carumbé | 353 bc | 66 ab | 418 ab |
|  | Logan | 290 ab | 47 a | 343 a |
|  | Baronesse | 230 a | 147 abc | 372 a |
|  |  | 356.4 *b* | 145.7 *a* | 504.6 *b* |
| Means | Prior | 441 | 203 | 646 |
|  | Ceibo | 464 | 157 | 624 |
|  | Quebracho | 304 | 179 | 491 |
|  | Danuta | 301 | 181 | 492 |
|  | Bowman | 241 | 141 | 393 |
|  | Carumbé | 289 | 90 | 388 |
|  | Logan | 244 | 29 | 335 |
|  | Baronesse | 184 | 109 | 299 |
|  |  | 308 | 136 | 458 |

**Table S9:** Type III ANOVA (Satterthwaite’s method) for phyllochron (Phy.) and FLN.

| Trait | Fixed effects | Sum Sq | Mean Sq | NDF | DDF | F value | Pr(>F) | % Variance |
| --- | --- | --- | --- | --- | --- | --- | --- | --- |
| FLN | Cultivar | 71.3 | 10.2 | 7 | 443.5 | 9.4 | *** | 7.65 |
|  | Year | 53.4 | 53.4 | 1 | 221.8 | 49.2 | *** | 5.74 |
|  | Sowing date (SD) | 702.4 | 702.4 | 1 | 444.6 | 646.9 | *** | **75.41** |
|  | Cultivar × Year | 16.8 | 2.4 | 7 | 443.3 | 2.2 | * | 1.8 |
|  | Cultivar × SD | 64.7 | 9.2 | 7 | 443.2 | 8.5 | *** | 6.95 |
|  | Year × SD | 18.4 | 18.4 | 1 | 443.9 | 16.9 | *** | 1.97 |
|  | Cultivar × Year × SD | 4.4 | 0.6 | 7 | 443.5 | 0.6 | 0.78 | 0.48 |
| Phy.  (e-a) | Cultivar | 32420 | 4631 | 7 | 398.5 | 28.21 | *** | 25.18 |
|  | Year | 16683 | 16683 | 1 | 90.6 | 101.60 | *** | 12.96 |
|  | Sowing date (SD) | 47486 | 47486 | 1 | 399.2 | 289.19 | *** | **36.88** |
|  | Cultivar × Year | 838 | 120 | 7 | 397.9 | 0.73 | 0.65 | 0.65 |
|  | Cultivar × SD | 14600 | 2086 | 7 | 397.5 | 12.70 | *** | 11.34 |
|  | Year × SD | 13719 | 13719 | 1 | 398.8 | 83.55 | *** | 10.65 |
|  | Cultivar × Year × SD | 3016 | 431 | 7 | 398.4 | 2.62 | * | 2.34 |

Significance codes: p < 0.01 ***; p < 0.01**; p < 0.05*.
